## Supplementary Figures 1 - 5; Supplementary Tables 1 - 6 for "Polar confinement of a macromolecular machine by an SRP-type GTPase"

**The file contains:**
Supplementary Figures 1 – 5
Supplementary Tables 1 – 6

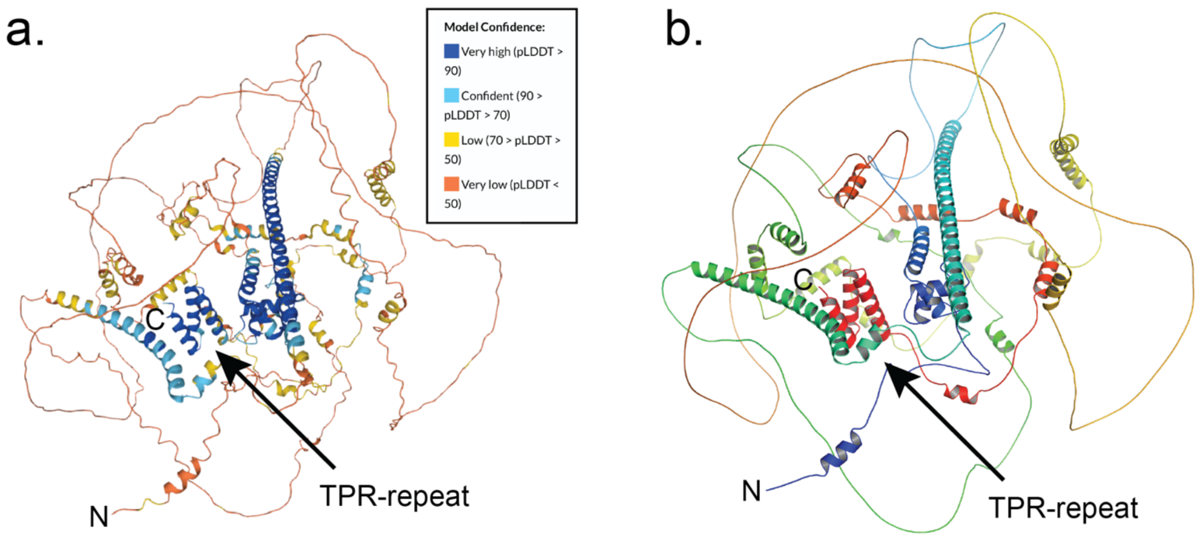


**Supplementary Figure 1. Alpha-Fold (1) model of HubP. a. Overall Confidence Assessment.** The confidence analysis of the HubP model reveals a predominantly low level of confidence. The accompanying inset provides a color-coded reference corresponding to the confidence scores. **b.** The Alpha-Fold model of HubP is presented in a simplified cartoon format, employing a rainbow color scheme to depict the sequence from N- to C-terminus. Notably, 'N' and 'C' signify the N-terminus and C-terminus, respectively.

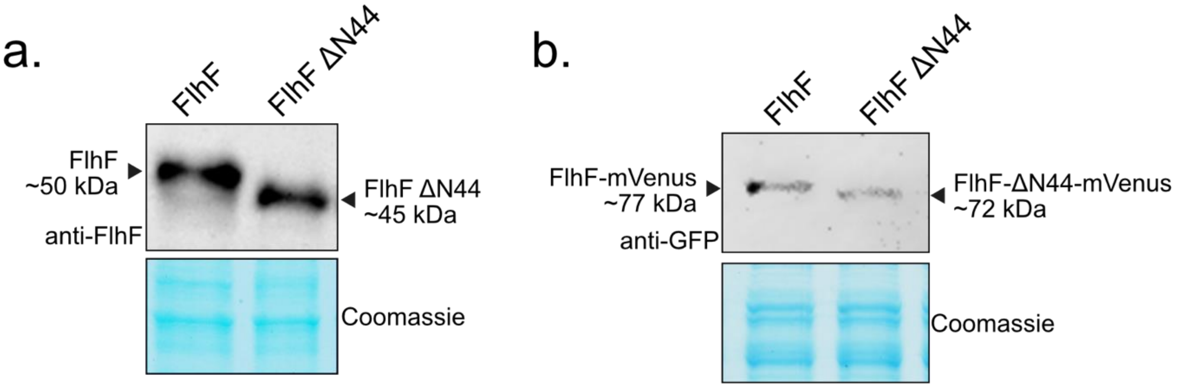


**Supplementary Figure 2. The deletion constructs of the 44 aa N-terminal domain of FlhF are stable produced in *Shewanella*.** Protein extracts from the respective *Shewanella* strains were separated by SDS-PAGE, transferred to membranes and the proteins were visualized by western blotting with antibodies against (**a.**) FlhF or (**b.**) GFP. For each strain the same OD units (10) were loaded for each sample.

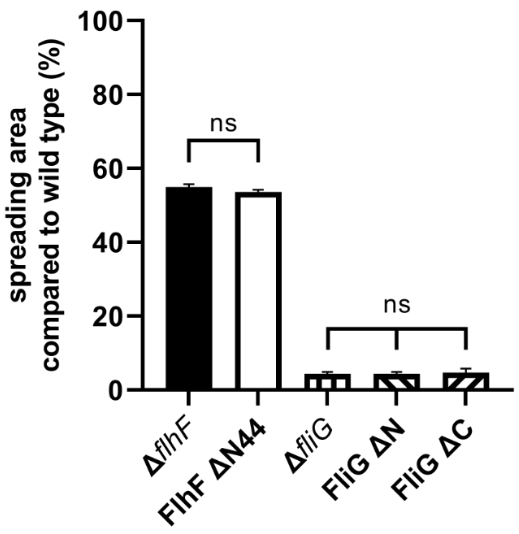


**Supplementary Figure 3. Spreading behavior of *S. putrefaciens* is impaired by deletion of the N- or C-terminal domain of FliG or the 44 aa N-terminal domain of FlhF.** Strains that are directly compared with each other were spotted on the same plate. Data from three independent experiments and the corresponding error bars are shown. A two-way ANOVA was performed (ns = not significant).

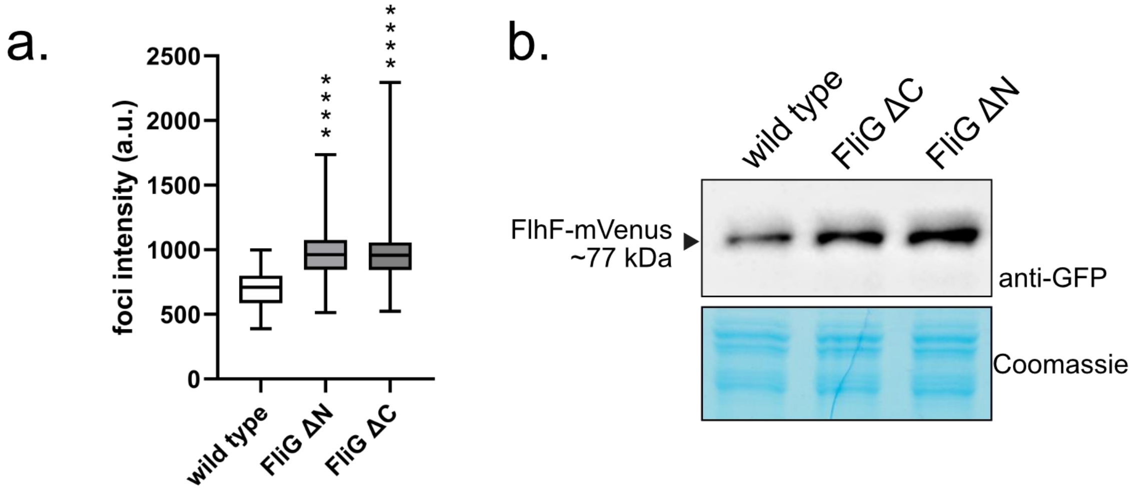


**Supplementary Figure 4. The deletion of the N- or C-terminal domain of FliG results in a FlhF-mVenus accumulation. a.** Quantification of FlhF-mVenus foci intensity (a.u. arbitrary units). Asterisks represent a p-value <0.0001 (Two-way ANOVA). **b.** Protein levels of FlhF-mVenus. Protein extracts from the respective *Shewanella* strains were separated by SDS-PAGE, transferred to membranes and the proteins were visualized by western blotting with antibodies against GFP.

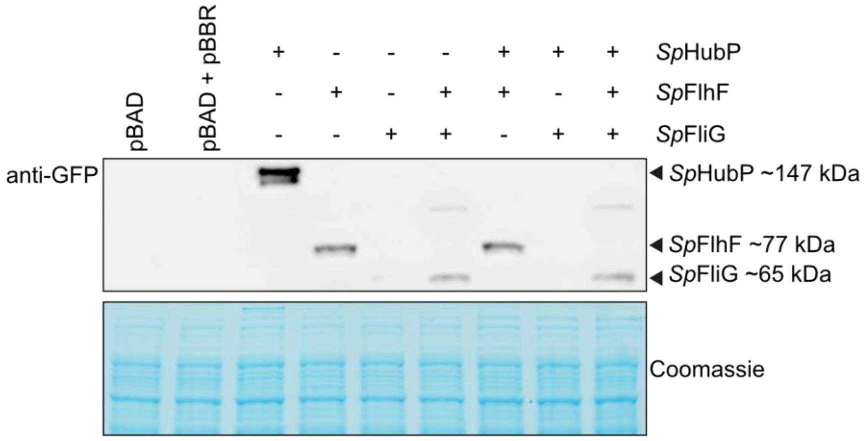


**Supplementary Figure 5. The *Shewanella* proteins are stable produced in *E. coli* DH5α.** Protein extracts from the respective *E. coli* strains were separated by SDS-PAGE, transferred to membranes and the *Shewanella* proteins HubP (*Sp*HubP), FlhF(*Sp*FlhF) or FliG (*Sp*FliG) were visualized by western blotting with antibodies against GFP. For each strain the same OD units (10) were loaded for each sample. pBAD and pBBR are loaded as empty vector controls.

**Supplementary Table S1.** Data collection and refinement statistics for the structure determination of the FlhF-FID domain.

| | |
|---|---|
| **Data collection** | |
| Space group | *P*$4_3$ |
| Cell dimensions | |
| *a*, *b*, *c* (Å) | 106.53 106.53 106.62 |
| α, β, γ (°) | 90 90 90 |
| Wavelength (Å) | 0.976990 |
| Resolution (Å) | 37.68 - 3.01 (3.118 - 3.01) |
| $R_{merge}$ | 0.07745 (2.6) |
| *I* / σ*I* | 19.39 (0.90) |
| Completeness (%) | 99.72 (99.83) |
| Redundancy | 13.5 (14.0) |
| *CC1/2* | 0.999 (0.485) |
| **Refinement** | |
| Resolution (Å) | 37.68 - 3.01 (3.118 - 3.01) |
| No. reflections | 23664 (2371) |
| $R_{work}$ / $R_{free}$ | 0.24/0.27 |
| No. atoms | 4269 |
| Protein | 4269 |
| Ligand/ion | 0 |
| Water | 0 |
| *B*-factors | 147.82 |
| R.m.s. deviations | |
| Bond lengths (Å) | 0.006 |
| Bond angles (°) | 0.76 |
| Ramachandran | |
| Favored (%) | 94.96 |
| Allowed (%) | 5.04 |
| Outliers (%) | 0.00 |

**Supplementary Table 2.** *Escherichia coli* strains used in this study.

| Strain | Genotype | Reference |
|---|---|---|
| **DH5αλ*pir*** | *sup E44, ΔlacU169 (ΦlacZΔM15), recA1, endA1, hsdR17, thi-1, gyrA96, relA1, λpirphage lysogen* | Miller & Mekanalos, 1988 |
| **WM3064** | *thrB1004 pro thi rpsL hsdS lacZ ΔM15 RP4-1360 Δ(araBAD) 567ΔdapA 1341: [erm pir(wt)]* | W. Metcalf, University of Illinois, Urbana-Champaign |

**Supplementary Table 3.** *Shewanella putrefaciens* CN-32 strains used in this study.

| Strain | Genotype | Reference |
|---|---|---|
| **wild type** | wild type strain of *S. putrefaciens* CN-32 | Fredricksonet al, 1998 |
| **Δ*flhF*** | deletion of the gene *flhF* (Sputcn32_2561) | (2) |
| **Δ*fliG*** | deletion of the gene *fliG* (Sputcn32_2575) | This study |
| ***flgE1* T183C** | markerless in-frame substitution of Thr183 to Cys in the polar hook protein FlgE1 (Sputcn32_2594) | (3) |
| ***flgE1* T183C Δ*flhF*** | markerless in-frame substitution of Thr183 to Cys in the polar hook protein FlgE1 (Sputcn32_2594), deletion of the gene *flhF* (Sputcn32_2561) | This study |
| ***flgE1* T183C FlhF ΔN44** | markerless in-frame substitution of Thr183 to Cys in the polar hook protein FlgE1 (Sputcn32_2594) and deletion of first N-terminal 44 amino acids of the gene flhF (Sputcn32_2561) | This study |
| **FlhF-1xGS-mVenus** | C-terminal mVenus tag of FlhF (Sputcn32_2561) linked with Gly and Ser | (4) |
| **FlhF ΔN44-1xGS-mVenus** | C-terminal mVenus tag of FlhF without first N-terminal 44 amino acids (Sputcn32_2561) linked with Gly and Ser | This study |
| **FlhF ΔN44** | Deletion of first N-terminal 44 amino acids of the gene *flhF* (Sputcn32_2561) | This study |
| ***flaB*$_1$ T166C *flaA*$_1$ T174C FliG$_1$ ΔN** | markerless in-frame substitution of Thr166 to Cys in the polar major flagellin protein FlaB1 (Sputcn32_2585) and Thr174 to Cys in the polar minor flagellin protein FlaA1 (Sputcn32_2586), deletion of the N-terminal domain (Δ2-85) of FliG(Sputcn32_2575) | This study |
| **FlhF-1xGS-mVenus FliG$_1$ ΔN** | C-terminal mVenus tag of FlhF (Sputcn32_2561) linked with Gly and Ser, deletion of the N-terminal domain (Δ2-85) of FliG(Sputcn32_2575) | This study |
| ***flaB*$_1$ T166C *flaA*$_1$ T174C FliG$_1$ ΔC** | markerless in-frame substitution of Thr166 to Cys in the polar major flagellin protein FlaB1 (Sputcn32_2585) and Thr174 to Cys in the polar minor flagellin protein FlaA1 (Sputcn32_2586),deletion of the C-terminal domain (Δ209-348) of FliG(Sputcn32_2575) | This study |
| **FlhF-1xGS-mVenus FliG$_1$ ΔC** | C-terminal mVenus tag of FlhF (Sputcn32_2561) linked with Gly and Ser, deletion of the C-terminal domain (Δ209-348) of FliG (Sputcn32_2575). | This study |

**Supplementary Table 4.** Plasmids used in this study.

**General plasmids:**

| Plasmid | Description | Reference |
|---|---|---|
| **pNPTS138-R6KT** | mobRP4+ ori-R6K *sacB;* β-galactosidase fragment alpha; suicide vector for *in-frame* deletions or integrations in *S. putrefaciens*; Kan$^r$ | (5) |
| **pBAD33** | Empty vector control | (5) |
| **pBBR1-MCS2** | Empty vector control | (5) |

***Escherichia coli* plasmids:**

| Plasmid | Description | Reference |
|---|---|---|
| **pBBR1-MCS2_HubP** | Expression of HubP (Sputcn32_2442) | This study |
| **pBAD33_HubP-GGS-sfGFP** | Expression of HubP (Sputcn32_2442) fused to GFP with Gly, Gly and Ser | This study |
| **pBAD33_FlhF_mVenus-GSGGG-FliG** | Expression of FlhF (Sputcn32_2561) and FliG (Sputcn32_2575) fused to mVenus with Gly, Ser, Gly, Gly, and Gly | This study |
| **pBAD33_mVenus-GSGGG-FliG** | Expression of FliG (Sputcn32_2575) fused to mVenuswith Gly, Ser, Gly, Gly, and Gly | This study |
| **pBAD33_FlhF-1xGS-mVenus** | Expression of FlhF (Sputcn32_2561) fused to mVenuswith Gly and Ser | This study |
| **pBAD33_mCherry-2xGGS-FliF_FliG** | Expression of FliG (Sputcn32_2575) and FliF (Sputcn32_2576) fused to mCherry with 2x Gly, Gly and Ser | This study |
| **pBAD33_FlhF_mCherry-2xGGS-FliF** | Expression of FlhF (Sputcn32_2561) and FliF (Sputcn32_2576) fused to mCherry with 2x Gly, Gly and Ser | This study |
| **pBAD33_FlhF_mCherry-2xGGS-FliF_FliG** | Expression of FlhF (Sputcn32_2561), FliG (Sputcn32_2575) and FliF (Sputcn32_2576)fused to mCherry with 2x Gly, Gly and Ser | This study |
| **pBAD33_mCherry-2xGGS-FliF** | Expression of FliF (Sputcn32_2576) fused to mCherrywith 2x Gly, Gly and Ser | This study |
| **pET24d_FliG1_N-His** | Expression of N-terminal His-tagged polar FliG | This study |
| **pET24d_FlhF_1-137_N-Strep** | Expression of N-terminal StrepII-tagged FlhF B-domain amino acids 1-137 | This study |
| **pET24d_FlhF_1-60_N-Strep** | Expression of N-terminal StrepII-tagged FlhF B-domain amino acids 1-60 | This study |
| **pET24d_FlhF_10-137_N-Strep** | Expression of N-terminal StrepII-tagged FlhF B-domain amino acids 10-137 | This study |
| **pET24d_FlhF_20-137_N-Strep** | Expression of N-terminal StrepII-tagged FlhF B-domain amino acids 20-137 | This study |
| **pET24d_FlhF_30-137_N-Strep** | Expression of N-terminal StrepII-tagged FlhF B-domain amino acids 30-137 | This study |
| **pET24d_FlhF_1-44_C-His** | Expression of C-terminal His-tagged FlhF B-domain amino acids 1-44 | This study |
| **pET24d_FlhF_C-Strep** | Expression of C-terminal StrepII-tagged FlhF | This study |
| **pET24d_FliG-MC_N-His** | Expression of N-terminal His-tagged polar FliG-MC domains | This study |
| **pET24d_FliG-NM_N-His** | Expression of N-terminal His-tagged polar FliG-NM domains | This study |

| | | |
|---|---|---|
| **pGAT3_FliF-C** | Expression of N-terminal GST-tagged polar FliF-C | This study |
| **pET24d_FliM_N-His** | Expression of N-terminal His-tagged polar FliM | This study |
| **pET24d_FliN** | Expression of polar FliN | This study |
| **pET24d_HubP-860-1097_N-Strep** | Expression of N-terminal StrepII-tagged HubP amino acids 860-1097 | This study |
| **pET24d_FlhF_C-His** | Expression of C-terminal His-tagged FlhF | This study |
| **pET24d_HubP-860-1033_N-Strep** | Expression of N-terminal StrepII-tagged HubP amino acids 860-1097 | This study |
| **pET24d_HubP-1033-1097_N-Strep** | Expression of N-terminal StrepII-tagged HubP amino acids 860-1097 | This study |
| **pET24d_FlhF-NG_C-His** | Expression of C-terminal His-tagged FlhF-NG | This study |
| **pET24d_FlhG_N-His** | Expression of N-terminal His-tagged FlhG | This study |

***Shewanella putrefaciens*** **CN-32 plasmids:**

| Plasmid | Description | Reference |
|---|---|---|
| **pNPTS138-R6KT FlhF-1xGS-Venus** | in frame complementation of *flhF* (Sputcn32_2561) with FlhF-1xGS-mVenus; Kan$^r$ | (4) |
| **pNPTS138-R6KT *flhF* KO** | deletion of the *flhF* gene (Sputcn32_2561); Kan$^r$ | (2) |
| **pNPTS138-R6KT *fliG* KO** | deletion of the *fliG* gene (Sputcn32_2575); Kan$^r$ | This study |
| **pNPTS138-R6KT FlhF ΔN44** | Deletion of first 44 N-terminal amino acids in *flhF* (Sputcn32_2561); Kan$^r$ | This study |
| **pNPTS138-R6KT FlhF ΔN44-GS-mVenus** | Deletion of first 44 N-terminal amino acids in *flhF*fused to mVenus (Sputcn32_2561); Kan$^r$ | This study |
| **pNPTS138-R6KT FliG ΔN** | deletion of the N-terminal domain (Δ2-85) of FliG(Sputcn32_2575); Kan$^r$ | This study |
| **pNPTS138-R6KT FliG ΔC** | deletion of the C-terminal domain (Δ209-348) of FliG (Sputcn32_2575); Kan$^r$ | This study |

**Yeast-Two Hybrid plasmids:**

| Plasmid | Description | Reference |
|---|---|---|
| **pGBKT7_FlhF** | pGBKT7 plasmid expressing the FlhF bait protein fused to the Gal4 DNA-binding domain | This study |
| **pGBKT7_FlhF-B** | pGBKT7 plasmid expressing the FlhF B-domain bait protein fused to the Gal4 DNA-binding domain | This study |
| **pGADT7_FliF-C** | pGADT7 plasmid expressing the FliFc prey protein fused to Gal4 activation domain | This study |
| **pGADT7_FliG_polar** | pGADT7 plasmid expressing the polar FliG prey protein fused to Gal4 activation domain | This study |
| **pGADT7_FliG_lateral** | pGADT7 plasmid expressing the lateral FliG prey protein fused to Gal4 activation domain | This study |
| **pGADT7_FlhG** | pGADT7 plasmid expressing the FlhG prey protein fused to Gal4 activation domain | This study |
| **pGADT7_FliM** | pGADT7 plasmid expressing the FliM prey protein fused to Gal4 activation domain | This study |
| **pGADT7_FliN** | pGADT7 plasmid expressing the FliN prey protein fused to Gal4 activation domain | This study |

| | | |
|---|---|---|
| **pGADT7_FlhF** | pGADT7 plasmid expressing the FlhF prey protein fused to Gal4 activation domain | This study |

**Supplementary Table 6.** Oligonucleotides used in this study. The abbreviations are: for: forward; rev: reverse; KO: knock-out (deletion) primer; KI: knock-in (integration) primer, CP: check primer, SP: sequencing primer, PC: plasmid construction

**Oligonucleotides <u>used for *Shewanella putrefaciens* CN-32 work:</u>**

| Name | Sequence | Purpose |
|---|---|---|
| **M13** | TGTAAAACGACGGCCAGTCC | CP/SP |
| **M13r** | CACACAGGAAACAGCTATGACC | CP/SP |
| **EcoRV-FlhFdN44-KO-for** | CAAGCTTCTCTGCAGGATAACGCGGCTAATTTCGCAGCA | KO |
| **OL-FlhFdN44-KO-rev** | GTAGAGGACGCATAAGTGGACTACGATGAGCCAAAAGCGAAGG | KO |
| **OL-FlhFdN44-KO-for** | TTTTGGCTCATCGTAGTCCACTTATGCGTCCTCTACTGGCCG | KO |
| **EcoRV-FlhFdN44-KO-rev** | GAATTCGTGGATCCAGATGGTAGTGTCGTAAAAGTGATGCAAAACCTATTA AATG | KO |
| **Check-FlhF-KO-for** | GCATCAGTCAATGCAAGCAACC | CP |
| **Check-FlhF-KO-rev** | GCACGGATTAATCCAGCATGCT | CP |
| **EcoRV-flhF KO-fwd** | GCCAAGCTTCTCTGCAGGATGCATAGGCGTCGGTGATTGAGG | KO |
| **OL-flhF KO-rev** | TAAGTGAAGGCATTTGAGTAGAGTTATGACCCTGG | KO |
| **OL-flhF KO-fwd** | CTCAAATGCCTTCACTTATGCGTCCTCTACTGG | KO |
| **EcoRV-flhF KO-rev** | GCGAATTCGTGGATCCAGATGCTAAGCATTCTCCTAAGCTTGTTG | KO |
| **PstI-fliG1-KO-fwd** | CCCTGCAGATGAAAATGGCCAAGTG | KO |
| **OL-fliG1-KO-rev** | TTAGAGGAACTCAGCCATTGTCTTGTAACCA | KO |
| **OL-fliG1-KO-fwd** | ATGGCTGAGTTCCTCTAATCCGAGTGACAATAA | KO |
| **PspOMI-fliG1-KO-rev** | TCCGGGCCCGATTCAATACCAAGACGTAAAGCG | KO |
| **Check-fliG1-KO-fwd** | ATTAGTGTTTCAGTTGCTGTGGAC | CP |
| **Check-fliG1-KO-rev** | AGGATAACGTCATCAGGGTGCA | CP |
| **EcoRV-FliG1 dN-fwd** | GCGAATTCGTGGATCCAGATGCTGCAGATGAAAATGGCCAAG | KO |

| | | |
|---|---|---|
| **OL-FliG1 dN-rev** | GTTAAAACCCATTCAGCCATTGTCTTGTAACCAG | KO |
| **OL-FliG1 dN-fwd** | GGCTGAATGGGTTTTAACAGTGAAGAATTCGTTCG | KO |
| **EcoRV-FliG1 dN-rev** | GCCAAGCTTCTCTGCAGGATCGGTCATCCACATCGATAAGGT | KO |
| **EcoRV-FliG1 dC-fwd** | GCGAATTCGTGGATCCAGATGCCTCTATCCTCAAGCACTTAG | KO |
| **OL-FliG1 dC-rev** | ACTCATTTATTTCGCTGCTTGTGCGCCACCT | KO |
| **OL-FliG1 dC-fwd** | GCAGCGAAATAAATGAGTTCCTCTAATCCGAGTGAC | KO |
| **EcoRV FLAG-FliG1 C rv** | GCCAAGCTTCTCTGCAGGATATTCAATACCAAGACGTAAAGCGG | KO |
| **Check-FliG1 dN-fwd** | GCAGTAGGATTCAGCACACAAC | CP |
| **Check-FliG1 dN-rev** | GCTTCTACCGTGACTGGTTTGA | CP |
| **EcoRV FlhF C-term fwd** | GCGAATTCGTGGATCCAGATGCAAGAAATGGTTGGACAGCCT | KI |
| **OL-FlhF-Venus rev** | CACGCTGCCCTCAAATGCACAGGCCATATTATCTG | KI |
| **OL_Venus fwd** | GCATTTGAGGGCAGCGTGAGCAAGGGCGAGGAGCTGTT | KI |
| **OL_Venus rev** | GTCATAACTTTACTTGTACAGCTCGTCCATGCC | KI |
| **OL-FlhF-Venus fwd** | TACAAGTAAAGTTATGACCCTGGATCAAGCAAG | KI |
| **EcoRV FlhF C-term rev** | GCCAAGCTTCTCTGCAGGATGCCACATCTAAAAATCGGTCGG | KI |

**Oligonucleotides used for *Escherichia coli* work:**

| Name | Sequence | Purpose |
|---|---|---|
| **pBAD33 for** | GCACGGCGTCACACTTTGCTATG | CP/SP |
| **pBAD33 rev** | GCCAGGCAAATTCTGTTTTATCAG | CP/SP |
| **pBBR1 for** | ACCATGATTACGCCAAGCGCG | CP/SP |
| **pBBR1 rev** | CGTTGTAAAACGACGGCCAGTG | CP/SP |
| **pBAD-OL-HubP-for** | AATTCGAGCTCGGTACCCAGGAGGGCAAATATGAAATTTCGCACTTCGTATCTTGTCGGT | PC |
| **OL-HubP-rev** | TCCTTTGCTGCTACCGCCACTAATCTCTTTTAGTAAACGTCCGGCCTC | PC |
| **HubP-gfp-for** | GAGATTAGTGGCGGTAGCAGCAAAGGAGAAGAACTTTTCACTGGAG | PC |
| **HubP-gfp-pBAD-rev** | CGACTCTAGAGGATCCCCTTAGGATCCTTTGTAGAGCTCATCCATGCC | PC |
| **Seq-HubP-1** | GCACAATCCGCTGATGTTGTTTC | SP |
| **Seq-HubP-2** | GATATTGCAGTAGCCGATACCGAG | SP |

| | | |
|---|---|---|
| **Seq-HubP-3** | ACGGGCTAAATCCAGTTTGGC | SP |
| **pBBR-HubP-for** | ATCGAATTCCTGCAGCCCAGGAGGGCAAATATGAAATTTCGCACTTCGTATCTTGTCGG | PC |
| **pBBR-HubP-rev** | AGAACTAGTGGATCCCCCTTAACTAATCTCTTTTAGTAAACGTCCGGCCTC | PC |
| **Seq-FlhF-1** | GGCTCAGATGCCGTTATCATGT | SP |
| **Seq-FlhF-2** | ATCGCATTGGCGCCTATGAGC | SP |
| **pBAD-OL-FlhF-for** | AATTCGAGCTCGGTACCCAGGAGGGCAAATATGAAGATTAAACGATTTTTTGCCAAAGACATGCG | PC |
| **OL-FlhF-rev** | GCCCTTGCTCACGCTGCCCTCAAATGCACAGGCCATATTATCTGACC | PC |
| **FlhF-mVenus-for** | TGTGCATTTGAGGGCAGCGTGAGCAAGGGCGAGGAGC | PC |
| **FlhF-mVenus-pBAD-rev** | CGACTCTAGAGGATCCCCTTACTTGTACAGCTCGTCCATGCCG | PC |
| **Seq-mVenus-1** | CGTCTATATCACCGCCGACAAGC | SP |
| **FlhF-FliG-rev** | CACCATATTTGCCCTCCTCTACTCAAATGCACAGGCCATATTATCTGAC | PC |
| **mVenus-FliG-for** | TAGAGGAGGGCAAATATGGTGAGCAAGGGCGAGGAGC | PC |
| **mVenus-FliG-rev** | CATGCCGCCGCCGCTGCCCTTGTACAGCTCGTCCATGCCG | PC |
| **FliG-linker-for** | AAGGGCAGCGGCGGCGGCATGGCTGAGAATAAAACAAAAGAAGCCGC | PC |
| **FliG-linker-rev** | CGACTCTAGAGGATCCCCTTAGAGGAACTCATCGCCACCACC | PC |
| **mVenus-FliG-for2** | AATTCGAGCTCGGTACCCAGGAGGGCAAATATGGTGAGCAAGGGCGAGGAGC | PC |
| **Seq-FliF-1** | GGCAATCTGCTTAGCCCTAGCC | SP |
| **Seq-FliF-2** | GGACTTTAAACCCGGTGCTGCA | SP |
| **Seq-FliF-3** | CTCTTGCTCAGTACGGGCAACG | SP |
| **Seq-mCherry-1** | GGGTTGGGAAGCAAGTTCAGAACG | SP |
| **mCherry-FliF-for** | AATTCGAGCTCGGTACCCAGGAGGGCAAATATGGTTTCCAAAGGGGAAGAGGACAATATGGC | PC |
| **mCherry-FliF-rev** | CGACTCTAGAGGATCCCCTCAGCCATTGTCTTGTAACCAGTTTTTCAC | PC |
| **FlhF-FliF-rev** | AACCATATTTGCCCTCCTCTACTCAAATGCACAGGCCATATTATCTGACC | PC |
| **FlhF-mCherry-FliF-for** | TAGAGGAGGGCAAATATGGTTTCCAAAGGGGAAGAGGACAATATGG | PC |
| **FliF-FliG-mCherry-rev** | GCTACCGCCGCTACCGCCTTTGTATAACTCATCCATACCACCAGTCGAATG | PC |
| **FliF-FliG-for** | GGCGGTAGCGGCGGTAGCATGATTGTCGGGTCTAATTCCGATTTAGCA | PC |

**Oligonucleotides used for Yeast-Two Hybrid constructs:**

| Name | Sequence | Purpose |
|---|---|---|
| **FlhF-BspHI-for** | TTAATCATGAGCGTGAAGATTAAACGATTTTTTGCCAAAGAC | PC |
| **FlhF-BamHI-rev** | TTAAGGATCCTTACTCAAATGCACAGGCC | PC |
| **FlhF-B-domain-BamHI-rev** | TTAAGGATCCTTACGGAATGTCTCGGCGCTCAG | PC |
| **FliF-C-NcoI-for** | TTAACCATGGGCCCAATGCTAAAACGCCTTATTTATCC | PC |
| **FliF-C-BamHI-rev** | TTTTGGCTCATCGTAGTCCACTTATGCGTCCTCTACTGGCCG | PC |
| **FliG1-PciI-for** | TTAAACATGTTGATGGCTGAGAATAAAACAAAAGAAGC | PC |
| **FliG1-BamHI-rev** | TTAAGGATCCTTAGAGGAACTCATCGCCACCAC | PC |
| **FliG2-NcoI-for** | TTAACCATGGGCGATAATTACGCCCAAGCAGC | PC |
| **FliG2-BamHI-rev** | TTAAGGATCCTTAGACAACGACCTGCTCTTC | PC |
| **FlhG-NcoI-for** | TTAACCATGGGAACCCTGGATCAAGC | PC |
| **FlhG-BglII-rev** | TTAAAGATCTTTATTCACTCGTTTTTTCCTC | PC |
| **FliM1-PciI-for** | TTAAACATGTGCGTGAGTGATTTATTAAGC | PC |
| **FliM1-BamHI-rev** | TTAAGGATCCTTACCCTGCATCCAACAACTC | PC |
| **FliN1-PciI-for** | TTAAACATGTAGCACAGAAGATACGGGC | PC |
| **FliN1-BamHI-rev** | TTAAGGATCCCGAATTAAAAAGCTTAAATAA | PC |

**Oligonucleotides used for protein expression constructs:**

| Name | Sequence | Purpose |
|---|---|---|
| **FliG-PciI-His-for** | TTAAACATGTTGCACCATCACCATCACCATATGGCTGAGAATAAAACAAAAG AAGCC | PC |
| **FliG-XhoI-rev** | TTAACTCGAGTTAGAGGAACTCATCGCCACCAC | PC |
| **FlhF-Strep-PciI-for** | TTAAGGTCTCCTCGAGTTATTTTTCGAACTGCGGGTGGCTCCAAGCGCTCCC CTCAAATGCACAGGCCATATTATCTGACC | PC |
| **FlhF-137-EcoRI-rev** | TTAAGAATTCTTACGGAATGTCTCGGCGCT | PC |
| **FlhF-60-EcoRI-rev** | TTAAGAATTCTTAAGGTGCAAGAGGTGTCGCGGAC | PC |
| **FlhF-ΔN10-Strep-PciI-for** | TTAAACATGTCTAGCTGGAGCCACCCGCAGTTCGAGAAAGGCTTAATTAACC ATATGCGTGCCGCTCTG | PC |
| **FlhF-ΔN20-Strep-PciI-for** | TTAAACATGTCTAGCTGGAGCCACCCGCAGTTCGAGAAAGGCTTAATTAACC ATACTCTCGGCTCAGATGCC | PC |
| **FlhF-ΔN30-Strep-PciI-for** | TTAAACATGTCTAGCTGGAGCCACCCGCAGTTCGAGAAAGGCTTAATTAACC ATAACAAAAAGGTGAATGGCGG | PC |
| **FlhF-NcoI-for** | TTAAGGTCTCCCATGGGCAAGATTAAACGATTTTTTGCCAAAGAC | PC |
| **FlhF-44-XhoI-rev** | TTAAGGTCTCCTCGAGAACCGCGGCGACAA | PC |
| **FlhF-Strep-XhoI-rev** | TTAAGGTCTCCTCGAGTTATTTTTCGAACTGCGGGTGGCTCCAAGCGCTCCC CTCAAATGCACAGGCCATATTATCTGACC | PC |

| | | |
|---|---|---|
| **FliG1-M-His-NcoI-for** | TTAACCATGGGCCACCATCACCATCACCATACGGCAGCATTAGGTG | PC |
| **FliG1-M-XhoI-rev** | TTAACTCGAGTTAGCCACCCATTTTCGCTGC | PC |
| **FliF-C-NcoI-for** | TTAACCATGGGCCCAATGCTAAAACGCCTTATTTATCC | PC |
| **FliF-C-His-XhoI-rev** | TTAACTCGAGTTAATGGTGATGGTGATGGTGGCCATTGTCTTGTAACCAG | PC |
| **FliM-His-NcoI-for** | TTAACCATGGGCCATCACCATCACCATCACGTGAGTGATTTATTAAGC | PC |
| **FliM-XhoI-rev** | TTAACTCGAGTTAAATACGATCTTGGGATG | PC |
| **FliN-PciI-for** | TTAAACATGTAGCACAGAAGATACGGGC | PC |
| **FliN-BamHI-rev** | TTAAGGATCCCGAATTAAAAAGCTTAAATAA | PC |
| **HubP-860-Strep-NcoI-for** | TTAACCATGGCTAGCTGGAGCCACCCGCAGTTCGAGAAAAGCGCGGTGCTG GATTGGGATACAGAG | PC |
| **HubP-XhoI-rev** | TTAACTCGAGTTAACTAATCTCTTTTAGTAAACGTCCG | PC |
| **FlhF-His-XhoI-rev** | TTAAGTCGACTTAGTGGTGATGGTGATGATGCTCAAATGCACAGGCCATATT ATC | PC |
| **HubP-1032-XhoI-rev** | TTAAGGTCTCCTCGAGTTAATCAATTATAGCGGCATCGC | PC |
| **HubP-1033-NcoI-for** | TTAAGGTCTCCCATGGGCGGTGCCTTACTTGGCG | PC |
| **FlhG-NcoI-for** | TTAACCATGGGCCACCATCACCATCACCATACCCTGGATCAAGCAAG | PC |
| **FlhG-XhoI-rev** | TTAACTCGAGTTATTCACTCGTTTTTTCCTCTT | PC |
